# AGENT: the Arabidopsis Gene Regulatory Network Tool for Exploring and Analyzing GRNs

**DOI:** 10.1101/2021.04.28.441830

**Authors:** Vincent Lau, Rachel Woo, Bruno Pereira, Asher Pasha, Eddi Esteban, Nicholas J. Provart

## Abstract

Gene regulatory networks (GRNs) are complex networks that capture multi-level regulatory events between one or more regulatory macromolecules, such as transcription factors (TFs), and their target genes. Advancements in screening technologies such as enhanced yeast-one-hybrid screens have allowed for high throughput determination of GRNs. However, visualization of GRNs in Arabidopsis has been limited to *ad hoc* networks and are not interactive. Here, we describe the Arabidopsis GEne Network Tool (AGENT) that houses curated GRNs and provides tools to visualize and explore them. AGENT features include expression overlays, subnetwork motif scanning, and network analysis. We show how to use AGENT’s multiple built-in tools to identify key genes that are involved in flowering and seed development along with identifying temporal multi-TF control of a key transporter in nitrate signaling. AGENT can be accessed at https://bar.utoronto.ca/AGENT.

## Introduction

Gene regulation is a complex process that encompasses the binding of often multiple transcription factors to the promoters of genes to induce expression these genes to help an organism respond to a particular environmental perturbation or to bring about a particular developmental program. Literature on *Arabidopsis thaliana* gene regulatory networks (GRNs) is rich and constantly growing. A large number of networks would be expected to exist, reflecting the complexity of regulation, and such networks would be active at different developmental stages and in response to environmental factors.

Regulatory networks also contain a diversity of modes of regulation, including protein-protein interactions (PPIs), protein-DNA interactions (PDIs) and miRNAs-mRNAs interactions (MMIs). In *Arabidopsis*, for PDIs alone, it is estimated that there are approximately 2,000 transcription factors (TFs), proteins that bind to DNA and regulate transcription (Wehner *et al*., 2011), and around 27,600 potential gene targets (Cheng *et al*., 2017). Clearly the potential number of networks and information in these networks has the potential to be very expansive. Consolidating this information into a database and providing tools to query it would help researchers understand which networks their genes of interest are involved in and would potentially uncover linkages between networks and genes.

Curating networks from the literature is essential to be able to notice large-scale patterns and to link data from different studies. While there are several examples of curated combined networks (Taylor-Teeples *et al*., 2015; Jin *et al*., 2015) they lack an interface to explore the data seamlessly. Along with annotation and data integration, information such as a gene’s expression level and subcellular location can easily be added to aid in hypothesis generation (Karlebach and Shamir, 2008). There are several examples of expression level association in the literature including Taylor-Teeples *et al*. (2015), who characterized how certain stresses, such as iron, salt, sulfur, and pH, would alter their secondary cell wall synthesis GRN by filtering the genes which were differentially expressed and subsequently investigating highly connected TFs that regulated those differentially expressed genes. Using this method, they confirmed their hypothesis that REVOLUTA (REV) was critical in lignification of secondary walls when the plant is under iron stress, as REV was shown to have many interactions with lignin biosynthesis genes. Other possibilities for integrating external data into networks can be found in review articles by Gaudinier & Brady (2016) and Liseron-Monfils & Ware (2015), who suggest integrating external PPI data to investigate additional types of regulation at the protein complex level. Last, GRNs can also be overlaid with functional data. Chen *et al*. (2019) constructed a yeast GRN to identify distinct subnetworks by their Gene Ontology (GO) categories to more easily reveal biological pathways defined by the subnetwork’s members. Clearly, the literature would seem to suggest that a web tool that integrates numerous types of information would be useful for hypothesis generation and network exploration.

Regulatory networks can be visualized and analyzed in by using graph theory notation. In this notation, edges are interactions between nodes, with nodes denoting genes, proteins, mRNAs or miRNAs. In doing so, GRNs can be analyzed with network theory to illuminate key regulatory areas (Alon, 2007). Such metrics include shortest path betweenness centrality (SPBC), which is based upon the shortest paths in a network, or degree centrality, which is the number of links incident on a node (Koschützki and Schreiber, 2008). SPBC involves counting the number of shortest paths a given node is involved in. This allows a researcher to estimate which nodes are essential for network communication and flow (Koschützki and Schreiber, 2008). Several papers in human and yeast models have noted that SPBC can be used in combination with other network statistics to identify significant genes (Bafna and Isaac, 2017; Sonawane *et al*., 2017).

Other significant nodes in networks can also be identified via network motifs or recurring patterns in regulation (Alon, 2007). A particular motif of note is the feed-forward loop in which one transcription factor (C) is jointly regulated by 2 others (A and B, where B is also regulated by A). The types of interactions in these motifs, i.e. activation or repression, can lead to different patterns of regulation. For instance it could be that the binding of *both* A *and* B to C is required to regulate C, or that the binding of *either* A *or* B to C is required to regulate C (Mangan and Alon, 2003). These patterns of regulation have led to them being of interest in many different gene regulatory network studies (Zhiponova *et al*., 2014; Sakuraba *et al*., 2015; X., Chen *et al*., 2019). In *Arabidopsis thaliana*, the walk-trap algorithm was used to identify highly connected nodes in leucine-rich repeat receptor kinases, illustrating that network analysis is a powerful analytic approach in plants (Smakowska-Luzan *et al*., 2018).

Current web tools for such networks exist but lack exploratory tools, the ability to integrate data, or usability. There is a dedicated PDI-visualizer tool, TF2Network (Kulkarni *et al*., 2018), however, the exploratory features are not expansive and the tool requires a gene list as input, which users may not always have. In contrast, ePlant (Waese *et al*., 2017), a tool that excels in external data integration, only shows interactions for a single transcription factor (TF) and not the entire GRN of which the TF is a member. AtRegNet, which is part of AGRIS (Yilmaz *et al*., 2011), a dedicated TF database with detailed summaries, appears to be in need of a big revamp to address aging web technology issues. Thus we present here AGENT, the Arabidopsis Gene Network Tool, available at https://bar.utoronto.ca/AGENT.

## Results

The AGENT interface is simple to navigate and has two parts, the landing page, where it is possible to enter gene names or AGI IDs, or to search for GRNs labeled with specific tags, such as Y1H (yeast one hybrid), see **Figure 1**. The results page (not shown) lists all GRNs that match the search criteria. Selecting a GRN then permits it to be explored and analyzed using tools included in AGENT, such as those to identify “Shortest Path” or “Find Selected Targets” (see lower panel in **Figure 1**). Options are also available to change the “Layout” or to analyze “Motifs”, as discussed later.

**Figure 1:**
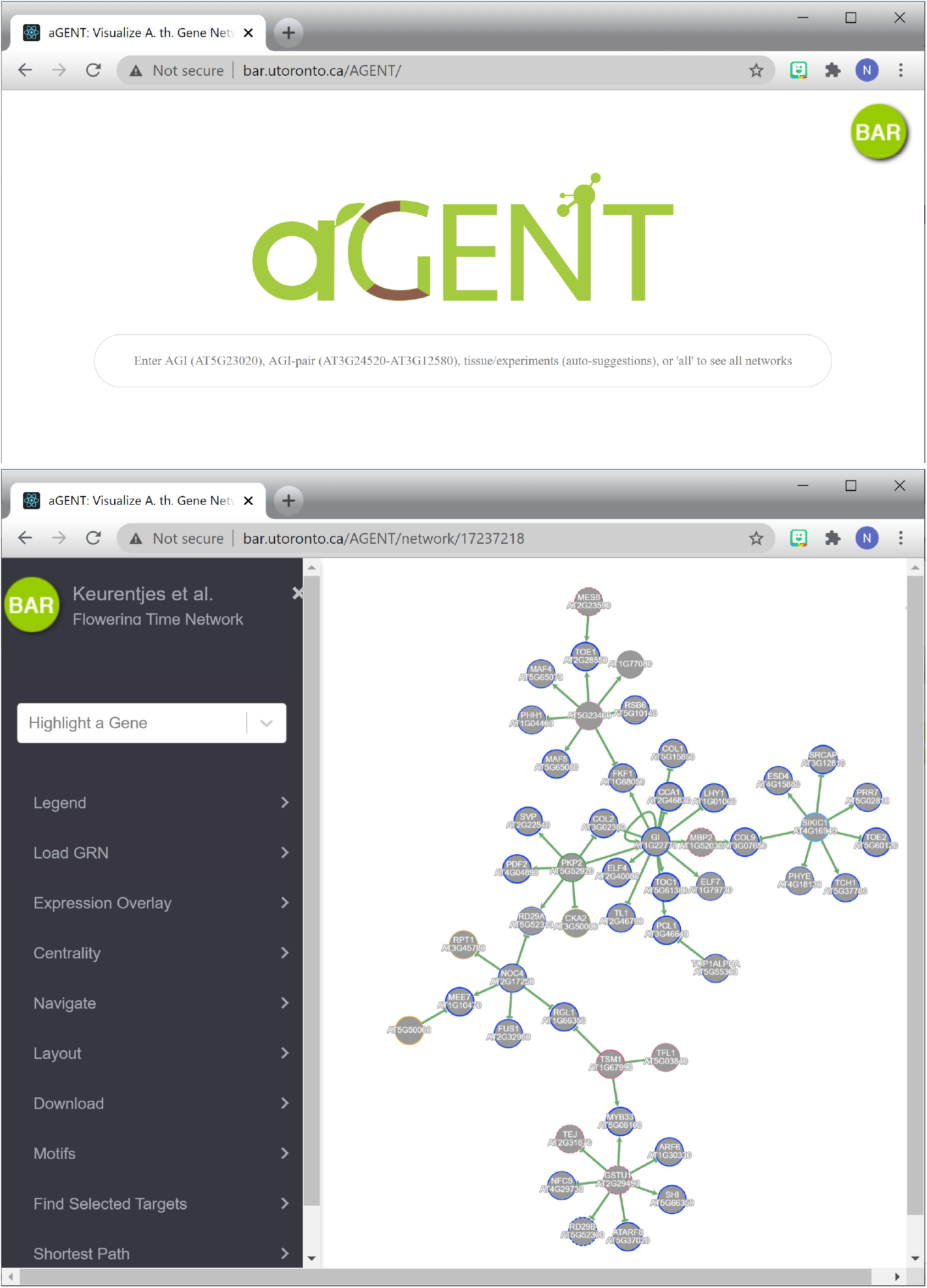
AGENT user interface. The top panel depicts the query page, while the bottom panel depicts the output page, showing the options available for display and analysis along the left side.

We have collected 13 Arabidopsis GRNs in AGENT, see **Table 1**. Rather than seeing this as “just” a useful curatorial exercise, one can see the utility of AGENT as a tool for exploration and analysis by examining the Keurentjes *et al*. (2007) flowering time network. In this example we will make use of several features of AGENT: “motif”, “betweenness centrality”, “Venn count” and “find selected targets”. When the feed-forward motifs from mfinder (Kashtan *et al*., 2004) are visualized for the network there are 3 genes/gene products of interest that create a coherent feed forward feedback loop: *GI* (*GIGANTEA*), *PKp2* (*PLASTIDIAL PYRUVATE KINASE 2*), and *ATCOL2* (*CONSTANS-like 2*), as shown in **Figure 2A**. This happens to be a “C3 OR” type feed forward loop (FFL), which effectively causes *ATCOL2* to be reduced in expression with a delay when either GI or Pkp2 increases in activity (Mangan and Alon, 2003). We can continue to explore this relationship by next adding betweenness centrality to the network. Now it becomes clear that the 2 high betweenness centrality genes/gene products, *PKp2* and *GI*, are receiving many input signals from the network to jointly regulate *ATCOL2*, indicative of potential crucial roles (Koschützki and Schreiber, 2008) for these genes/gene products in the flowering time network, which was in part derived from eQTLs.

**Figure 1:**
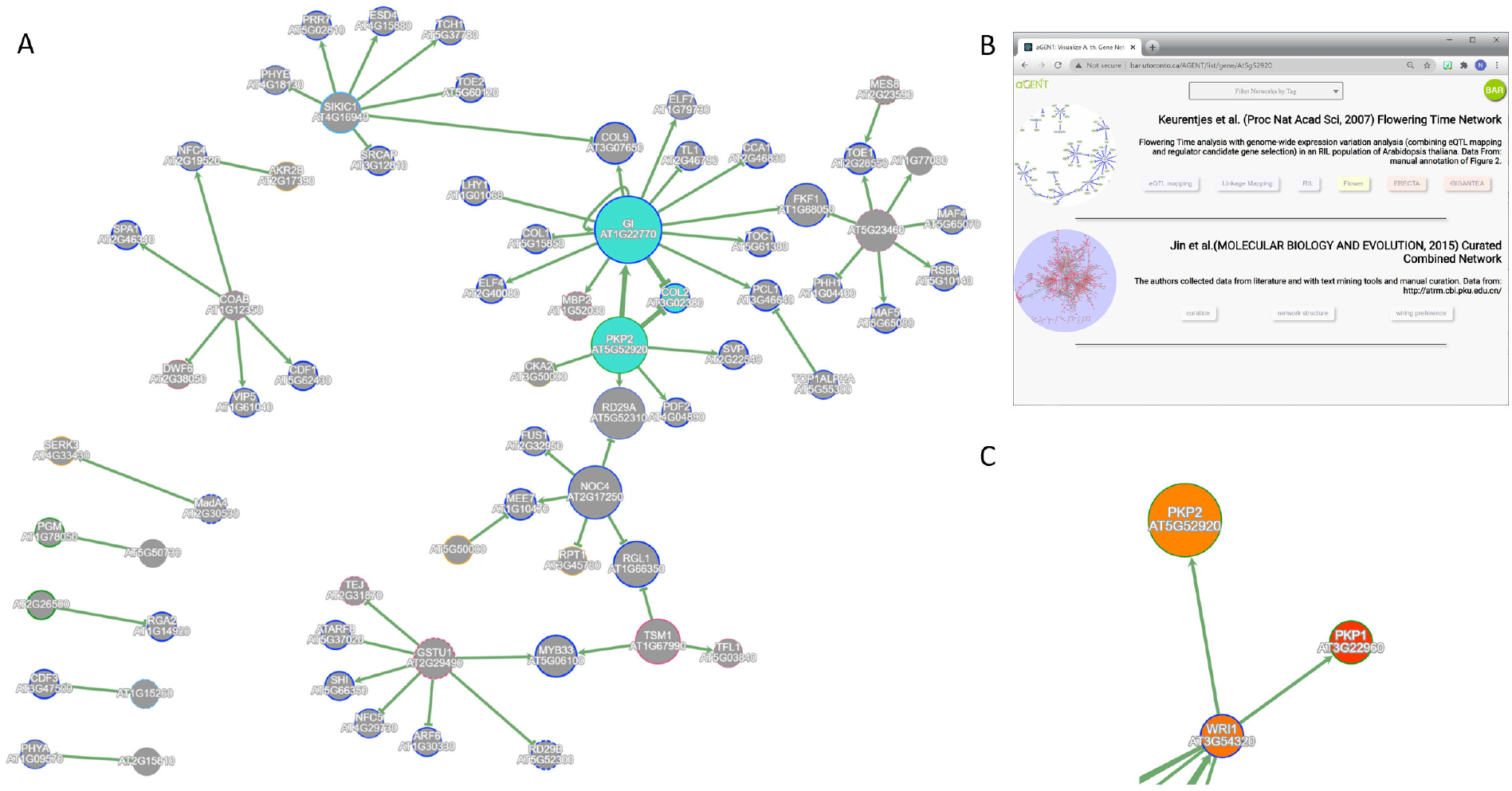
Using AGENT to identify key genes and time points of network activity. **A**. flowering time network from Keurentjes et al. (2007), with a feed forward loop (bold lines between nodes in cyan) for three nodes with high betweenness centrality (denoted by node size). **B**. AGENT search results for Pkp2 (At5g52920), showing two networks with this gene/gene product. **C**. WRI1 regulates *PKP2* expression via a protein-DNA interaction and is expressed at a high expression level in maturing seeds (red = 100% of expression potential).

**Table 1:**
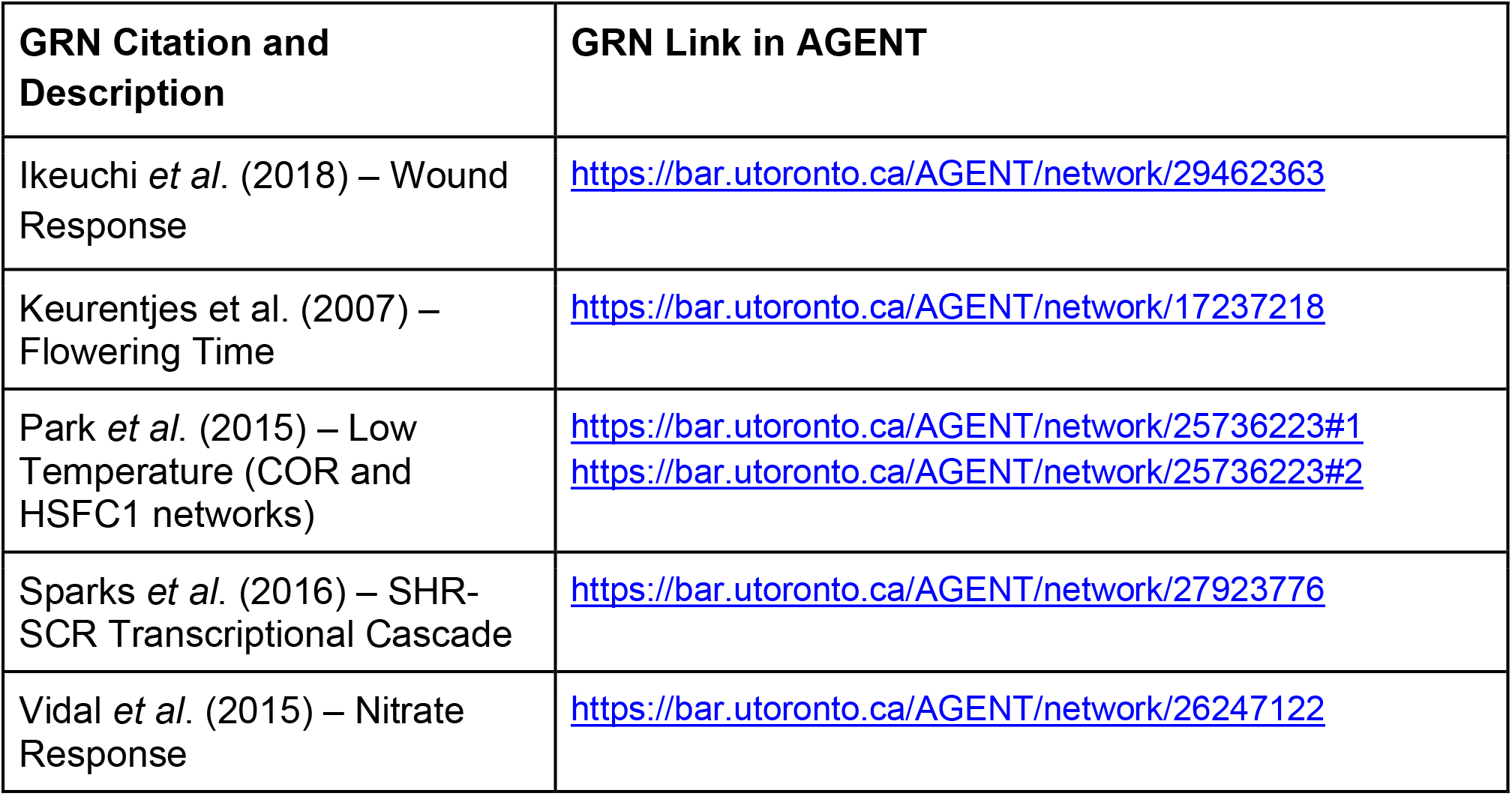
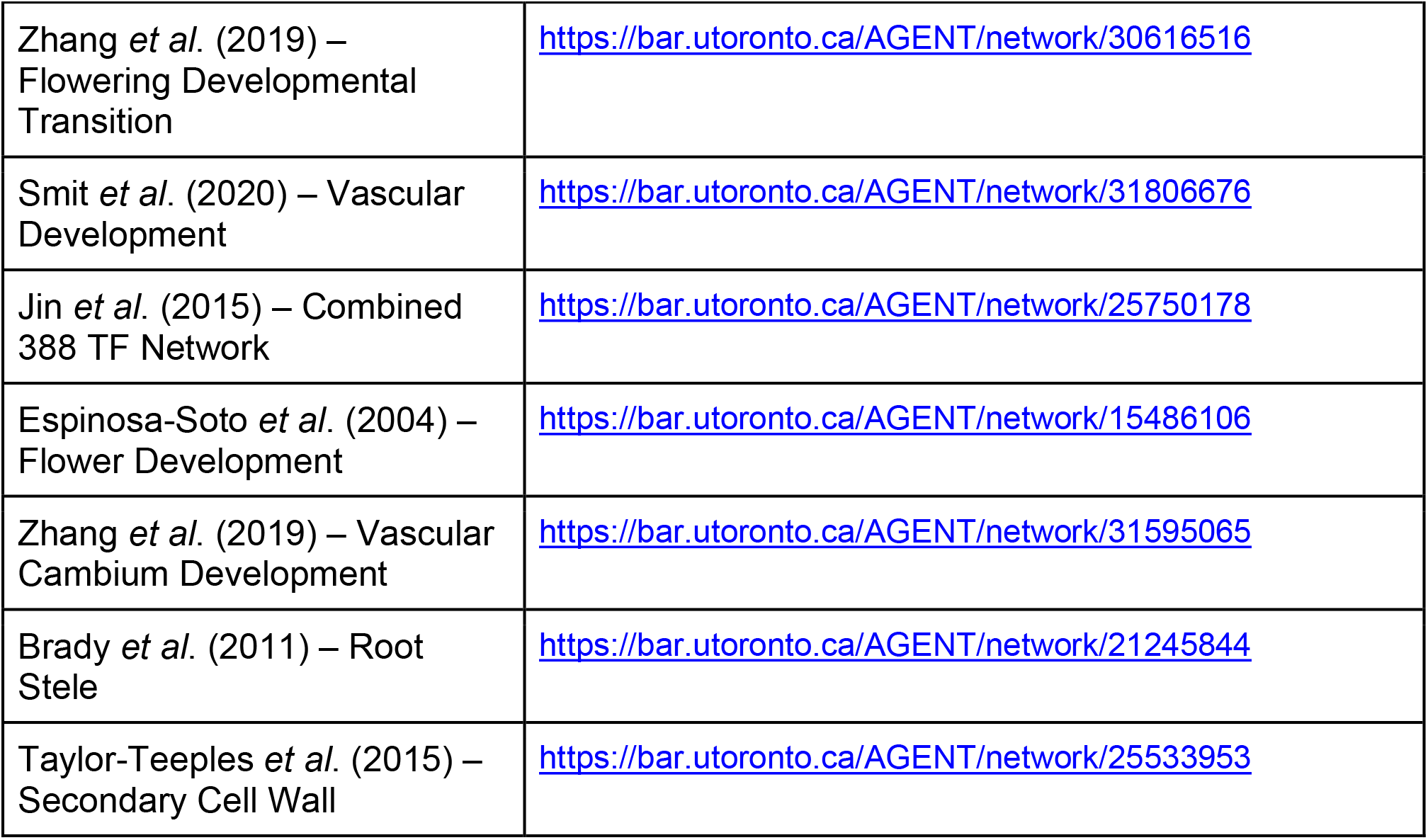
Manually Annotated Networks and their AGENT links.

| **GRN Citation and Description** | **GRN Link in AGENT** |
| --- | --- |
| Ikeuchi *et al.* (2018) – Wound Response | https://bar.utoronto.ca/AGENT/network/29462363 |
| Keurentjes *et al.* (2007) – Flowering Time | https://bar.utoronto.ca/AGENT/network/17237218 |
| Park *et al.* (2015) – Low Temperature (COR and HSFC1 networks) | https://bar.utoronto.ca/AGENT/network/25736223#1<br>https://bar.utoronto.ca/AGENT/network/25736223#2 |
| Sparks *et al.* (2016) – SHR-SCR Transcriptional Cascade | https://bar.utoronto.ca/AGENT/network/27923776 |
| Vidal *et al.* (2015) – Nitrate Response | https://bar.utoronto.ca/AGENT/network/26247122 |
| Zhang *et al.* (2019) – Flowering Developmental Transition | https://bar.utoronto.ca/AGENT/network/30616516 |
| Smit *et al.* (2020) – Vascular Development | https://bar.utoronto.ca/AGENT/network/31806676 |
| Jin *et al.* (2015) – Combined 388 TF Network | https://bar.utoronto.ca/AGENT/network/25750178 |
| Espinosa-Soto *et al.* (2004) – Flower Development | https://bar.utoronto.ca/AGENT/network/15486106 |
| Zhang *et al.* (2019) – Vascular Cambium Development | https://bar.utoronto.ca/AGENT/network/31595065 |
| Brady *et al.* (2011) – Root Stele | https://bar.utoronto.ca/AGENT/network/21245844 |
| Taylor-Teeples *et al.* (2015) – Secondary Cell Wall | https://bar.utoronto.ca/AGENT/network/25533953 |

In order to get a sense of where these genes might play additional roles, we can identify networks containing the genes in the above motif, by searching for them on the AGENT landing page. **Figure 2B** contains the results of searching for *Pkp2* (AT5G52920). In this case, the Jin et al. (2015) “Curated Combined” network encompassing 388 transcription factors and their targets also contains *Pkp2*. When we load this GRN we can note the following interaction: “AT3G54320 pdi-a AT5G52920” (AT3G54320 is *WRINKLED1* gene and the *pdi-a* terminology in AGENT denotes a protein-DNA interaction with the ability to activate expression), which supports a statement by Baud et al. (2007), “together with other genes encoding enzymes involved in glycolysis and fatty acid biosynthesis, *PKp1* and *PKp2* may consequently be targets of WRI1 […] The *pkp* phenotypes described in this study are fully consistent with this regulatory model that emphasizes the key role of plastidial metabolism, and its tight control in early-maturing seeds of *A. thaliana*”. We can test the hypothesis that both *Pkp2* and *PKp1* are expressed during seed development at expression levels near the maximum for all these genes in maturing seeds: **Figure 2C** shows the high expression potential of these two genes in seeds at the walking-stick stage of embryo development. *WRI1* is also at a high expression potential in developing seeds, again see **Figure 2C**. Interestingly, overexpression of *WRI1* in *Brassica napus* accelerates flowering (Li *et al*., 2015), and thus we could hypothesize from the Keurentjes et al. (2007) network that the link to flowering time could be through Pkp2.

We can also use AGENT’s motif scanner tool to quickly identify important TFs that regulate downstream genes in a particular GRN. For example, in the summarized Vidal et al. (2015) nitrate signaling network (see **Figure 3A**), we can quickly identify that there are 6 motifs which are FFLs.

**Figure 3.**
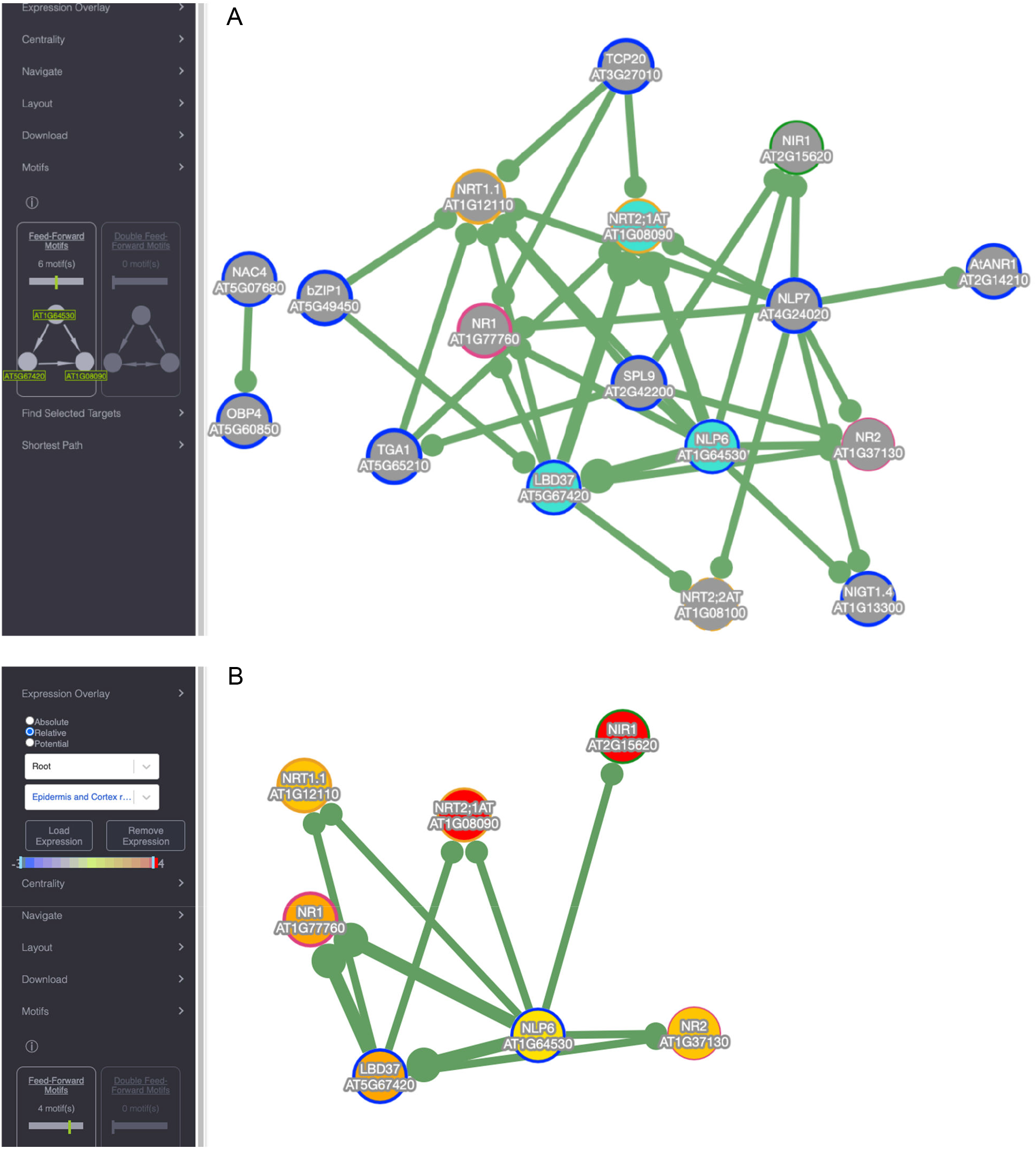
Identifying feed forward motifs in the Vidal et al. (2015) nitrate signaling network using AGENT’s Motifs tool. **A**. One of 6 feed forward motifs in the networks, highlight by the cyan nodes. **B**. Immediate neighbours of NLP6 constitute 4 feed forward motifs. Nitrate response gene expression profiles (as fold change relative to low nitrogen controls) in epidermis and cortex cells are overlaid on the nodes, with red being log_2_ ~3.4 increase (~11-fold) in expression relative to the control.

In particular, 4 of the 6 FFLs identified contain the TF NLP6 (AT1G64530), which regulates another TF gene, *LBD37* (AT5G67420), and, in turn, both regulate downstream target genes, such as *NRT2* (AT1G08090). A researcher can select NLP6 via the right-click menu to only display its immediate neighbours to simplify the network and focus on the 4 FFLs (see **Figure 3B**). NLP6 has been suggested to activate genes containing nitrate-responsive *cis*-elements (NREs) in their promoters via post-translational modifications (PTMs) near the N-terminus of NLP6 rather than via nitrogen-mediated induction of expression of the *NLP6* gene (Konishi and Yanagisawa, 2013). In contrast, its downstream target and FFL coactivator, *LBD37*/LBD37, has been shown to be strongly induced by nitrate and represses nitrate-responsive genes such as nitrate transporter (*NRT*) genes upon induction (Zhao *et al*., 2018). Thus, NLP6 activates *NRT2* and *LBD37*, while LBD37 represses *NRT2* after activation thus forming an incoherent type-1 FFL (I1-FFL), which is one of the most common FFLs (Alon, 2007). I1-FFLs are believed to act as response accelerators by allowing a rapid rise in the dually targeted gene (in this case *NRT2*) before being repressed due to subsequent repressor induction. Response accelerators have experimentally been shown to be useful in nutrient uptake, such as sugar uptake in *E. coli*. To see whether *NLP6* is not induced by nitrate while *LBD37* is induced by nitrate, nitrate root (epidermis/cortex) treatment expression profiles from Gifford et al. (2008) were overlaid in the AGENT interface (see **Figure 3B**). We see that *NLP6* expression is only modestly increased (log_2_ induction = 1.2, around a 2.3-fold increase) as expected, while *LBD37* is strongly induced (log_2_ induction = 2.8, a ~5-fold increase), likely due to *NLP6* activation. Their target gene, *NRT2* is also highly expressed (log_2_ induction = 3.4, an ~11-fold increase). Since the roots were nitrogen-depleted and expression-profiling was done shortly thereafter (2h), it is likely that this I1-FFL allows a rapid uptake of nitrate by highly expressing *NRT2* during nitrate treatment before being attenuated by LBD37. Indeed, Rubin et al. (2009), Scheible et al. (2004), and Zhao et al. (2018) saw similar results with *NRT2* being highly expressed after 3h, 3h, and 2h nitrate treatment, respectively. Moreover, Cerezo et al. (2001) showed that nitrate influx and *NRT2* expression strongly peaks 6-12 hours after nitrate treatment before returning to baseline levels in wildtype roots as expected by the I1-FFL model, whereas *nrt2* mutants show a flattened response. To conclude, by using AGENT’s Motifs tool and *ad hoc* knowledge regarding TFs and combining this with expression data, researchers can investigate the relationships between multiple TFs and target genes, particularly their roles in temporal regulation.

## Discussion

We describe the creation of a web tool for exploring gene regulatory networks in Arabidopsis thaliana, AGENT. It can be used to answer simple questions such as if one’s gene of interest is found in a particular GRN. But coupled with a researcher’s own biological knowledge more sophisticated analyses are also possible. For example, by using AGENT, a researcher can identify feed forward motifs and overlay expression information from the BAR’s extensive collection of gene expression data to identify biological conditions in which a GRN might be active. We will continue to curate Arabidopsis GRNs and encourage submissions from the community as SIF files.

## Materials and Methods

### GRN Curation and Annotation

GRNs included in the manual curation process were found in *Arabidopsis thaliana* and came from both from predictive algorithms and experimental methods. After being identified as a GRN of interest, the GRN data were manually curated into SIF (Simple Interactions Format) files with tags relating to Experiment, Condition, Genes, and a miscellaneous category. The data in these files was loaded into a MySQL database hosted on the BAR (Bio-Analytic Resource for Plant Biology).

### Application Development

To serve AGENT, REST (Representational State Transfer) API (Application Programming Interface) development was done with Express.js (Node.js; https://nodejs.org) to connect to the aforementioned MySQL database. This REST API returns interaction data along with information pertaining to each curated network. For example, the API endpoint https://bar.utoronto.ca/interactions_api/tags/Y1H retrieves all GRNs that used the Yeast-One-Hybrid technique. API development was also used for AGENT’s motif scanning tool where we utilized mFinder 1.21 (Mangan *et al*., 2003) with the following parameters (directed, r = 100, s = 3, u = 4, z = 2) to quickly identify FFLs. Parameters were recommended by the authors and optimized for quick web retrieval. AGENT also uses additional APIs we host at the BAR such as SUBA4 (SUBcellular localization database for Arabidopsis proteins; Hooper *et al*., 2017) localization data, ePlant expression data, interactions data, and gene annotations.

As part of this effort, we also created a database describing the highest levels of expression for each gene for all of the compendia accessed by ePlant (Waese *et al*., 2017) and the eFP Browser (Winter *et al*., 2007). We use this database, which captures in which samples/compendia the top 10 highest levels of expression for a given gene occur, in order to be able to compute for the genes in the AGENT interface how strongly they are expressed relative to their potential maximum expression, expressed as a percent. Expression may also be viewed as an absolute value, or as fold-change relative to the appropriate control.

We then developed the web-based user-interface (UI) with React (https://reactjs.org) along with React-Router to allow pagination and ePlant integration. Cytoscape.js (Franz *et al*., 2016) along with additional UI plugins (such as cytoscape-popper which enables tooltips) was used to visualize the networks. Cytoscape.js was also used to perform network analysis (e.g., degree centrality). User-testing was performed by webcasting a user’s experience with AGENT via open exploration and tasked objectives. Source code for the AGENT UI can be accessed via https://github.com/VinLau/aGENT.

## Acknowledgements, Contributions and Funding

We are grateful to Max Franz for helpful Cytoscape.js discussions. VL and RW created the AGENT user interface. VL and AP created the SQL database and backend APIs. BP developed the database for use in calculating expression potential. EE worked on the AGENT database. NJP, RW, and VL wrote the manuscript. NJP and VL created the figures and edited the manuscript. All authors have approved the manuscript. NJP was funded by Genome Canada/Ontario Genomics grant OGI-162.

## Notes

http://bar.utoronto.ca/AGENT

### Competing Interest Statement

The authors have declared no competing interest.

